# A Unified Deep Learning-Based Framework for Reference-Based and Reference-Free Local Ancestry Inference

**DOI:** 10.64898/2026.08.01.742246

**Authors:** Thomas Diem

## Abstract

Local ancestry inference (LAI) identifies the ancestral origin of genomic segments within admixed individuals and is an important tool for population genetics and disease association studies. Existing LAI methods rely on reference panels composed of individuals from ancestral populations, limiting their applicability when such panels are unavailable or poorly characterized. We present Optional Reference Inference Ancestry Network (ORIAN), a software package containing two complementary algorithms for local ancestry inference. The first is a reference-based approach that combines neural network predictions with a hidden Markov model to produce probabilistic ancestry assignments. The second is a reference-free method that introduces an iterative framework in which admixed individuals are used as probabilistic references for one another, enabling local ancestry inference without labeled ancestral reference panels. Both methods are trained on a diverse set of simulated admixture scenarios to promote generalization across populations. We evaluate ORIAN on human, *Drosophila melanogaster*, and fully simulated datasets, comparing its performance against RFMix and LOTER across a range of admixture times and proportions. In the reference-based setting, ORIAN achieves the highest median diploid accuracy for recent admixture while remaining competitive across a broad range of scenarios. In the reference-free setting, ORIAN produces competitive local ancestry estimates using only admixed individuals, extending local ancestry inference to settings where ancestral reference panels are unavailable. These results demonstrate that ORIAN provides an accurate and flexible framework for both conventional and reference-free local ancestry inference.

## 1 Introduction

Local ancestry inference (LAI) is the task of determining the ancestral origin of individual segments of an admixed genome [1, 2]. It plays an important role in understanding population history, detecting selection, and improving the resolution of association studies in diverse populations [3, 4]. Most existing approaches to LAI rely on the availability of reference panels consisting of individuals from the ancestral source populations, including methods such as RFMix and ELAI [2, 5]. However, such panels may not always exist, may be imperfectly matched to the true sources, or may be too small to yield reliable results.

These limitations motivate the development of local ancestry methods that do not rely on reference panels. Inferring local ancestry in this setting is significantly more challenging, as the ancestral populations must be inferred jointly with the ancestry of each sample [6, 7]. Nevertheless, many applications, including those involving ancient DNA, isolated populations, or poorly studied species, require such capabilities [8, 9].

In this work, we introduce the software package Optional Reference Inference Ancestry Network (ORIAN), which contains two complementary algorithms for local ancestry inference, both based on a shared neural network architecture. The first is a reference-based method that combines neural network outputs with a sequential HMM-based framework to produce accurate, probabilistic ancestry assignments. The second is a reference-free approach that uses a novel iterative scheme in which admixed individuals are treated as probabilistic references for one another. Both algorithms are trained on a wide range of simulated admixture scenarios to ensure generalizability across datasets. To our knowledge, ORIAN is the first method to extend accurate local ancestry inference to settings in which only admixed individuals are observed.

We evaluate our method on three datasets — human genomes, *Drosophila melanogaster*, and fully simulated populations — and show that the reference-based version outperforms existing methods across a range of admixture times and proportions. Furthermore, our reference-free method achieves comparable accuracy to leading reference-based methods, despite not using any known ancestry labels. Together, these results demonstrate the flexibility and robustness of our approach in both standard and challenging settings.

## 2 Methods

### 2.1 Training Pipeline

#### 2.1.1 Generating data for model training

To capture a wide range of biologically relevant scenarios, we developed a pipeline that combines the coalescent simulator MACS [10] to model population divergence with SELAM [11] to simulate ancestry switch points in admixed individuals. Key parameters, including admixture proportion, admixture time, ancestral population size, divergence time, daughter population sizes, recombination rate, mutation rate, and chromosome length, are sampled randomly and independently within ranges reflecting plausible biological events. For model training, we generate 400 simulations and reserve 50 simulations for validation, with each simulation containing a substantial number of both reference and admixed individuals.

#### 2.1.2 Neural network architecture

To fully capture the structure of each input example, we design a neural network that accepts a four-dimensional tensor as input. The architecture integrates position-independent transformer attention mechanisms with nonlinear operations and normalization steps applied across multiple tensor modes. Specifically, each input tensor has shape (3, 48, 10001, 38) and is used to predict the ancestry of a single SNP in an admixed individual. The first mode of length 3 represents possible classifications: homozygous for either ancestry or heterozygous. Each subtensor along this mode is processed through the network and reduced to a single output representing the class probability, with normalization intermittently applied across this mode.

The second and third modes, of length 48 and 10,001, correspond to the maximum number of reference individuals and SNPs included in a forward pass, respectively. The final mode of length 38 encodes SNP-specific information, including the alleles of the reference and admixed individual, the label of the reference individual, and the potential label of the admixed individual (indexed along mode 1). This results in 3^4^ = 81 possible encodings, which can be reduced to 36 by exploiting invariances between the two homozygous allele types and between the two homozygous ancestries. Each encoding is represented as a one-hot vector.

In addition, probabilities of a switch point between each SNP and the SNP of interest—conditioned on both starting ancestries—are computed using admixture time and admixture proportion, and concatenated to the input along the last dimension. Positions or individuals without information are encoded as a zero vector. This occurs when there are fewer than 48 reference individuals, when the probability of a switch point between the middle SNP and the SNP of interest is below a specified bound, or when the SNP of interest is near the sequence boundaries. The first two cases also enable computational efficiency during inference, as the learned parameters can be transferred to a smaller model at the start of the inference algorithm.

#### 2.1.3 Training Process

For the scenario with a reference panel, each training example is constructed by randomly selecting one admixed individual. Reference panel sizes for each class are drawn from a hypergeometric distribution, and reference individuals, the admixed individual, and the SNP to be predicted are all chosen randomly. Invariant sites are pruned on the fly based on the subset of individuals selected from the simulation. This process is repeated for every training simulation in each epoch, and the model is trained for approximately 3,000 epochs using the Adam optimizer with Adaptive Nesterov Momentum and exponential learning rate decay.

For the scenario without reference panels, training occurs in two phases. In the first phase, the process mirrors that of the reference-panel model, except that other admixed individuals are used as reference individuals. In the second phase, the model is trained to generalize to probabilistic labels and differently structured examples that would be present during inference. To achieve this, we generate training examples directly from the inference algorithm: we run the algorithm on many simulated examples, randomly save inputs and their correct outputs from forward passes, and then train the model on these saved examples using a low learning rate until overfitting begins. Because updates to the neural network parameters alter the types of examples encountered during inference, this process is performed iteratively. Starting from the phase-one model, the model is updated with a new dataset at each iteration. Iterations continue until negligible improvement is observed, which required about 20 iterations in practice.

### 2.2 Inference algorithm

#### 2.2.1 Neural network incorporation

Neural network inference forms the core of both algorithms. When a reference panel is sufficiently large or exhibits strong class imbalance, inference is divided into multiple forward passes. These passes are organized to maximize the number of reference individuals per pass, maintain roughly equal pass sizes, and minimize class imbalance. To achieve this, individuals may be reused, provided that each individual from a given ancestry appears in the same number of forward passes. Invariant sites are pruned at the start of each pass, and the outputs of multiple passes are combined using a weighted average.

Similarly, in scenarios without a reference panel, if there is a large number of admixed individuals, inference at each iteration is split into multiple forward passes. The groups of individuals are randomized at each timestep, and invariant sites are pruned on the fly.

#### 2.2.2 Optional fine-tuning on reference panel

As the neural network is pretrained on a wide variety of simulations, we include an optional fine-tuning stage at the start of the algorithm to further adapt the model using data from the reference panel. To generate training examples, we simulate admixed individuals from the reference panel based on specified admixture proportions and times. Each simulated individual is constructed by randomly sampling haplotype segments from reference haplotypes according to simulated switch points. Reference individuals are generated by randomly combining haploid sequences. Although this step requires computational phasing, the procedure is generally robust to phasing errors, as the haploid sequences are only used for simulation purposes. Training is fully automated, with a ReduceLROnPlateau scheduler adjusting the learning rate. The model parameters achieving the best test set performance replace the original parameters and are subsequently used for inference.

#### 2.2.3 With reference panels: combining results via an HMM-based solution

After the neural network infers ancestry at a fixed set of SNPs along an admixed chromosome, these outputs must be combined to produce a full chromosomal solution. Given the sequential nature of the outputs and prior knowledge of transition probabilities, a Hidden Markov Model (HMM) approach follows naturally from Bayes’ theorem. Switch points along a chromosome can be estimated by a Poisson process where we consider tracts to be independent. We construct the prior transition matrix as

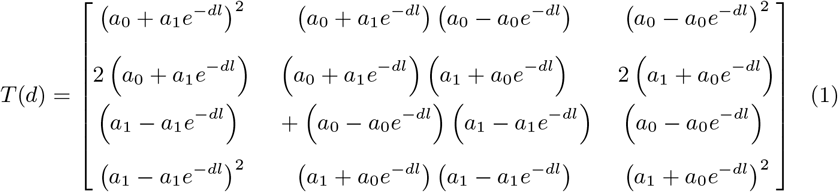

as a function of genetic distance *d* where

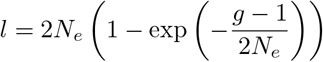

for *g* generations since admixture, effective population size *N*_*e*_, and admixture fraction *a*_0_ for population 0 and *a*_1_ = 1 − *a*_0_ for population 1 [12]. The classes represent homozygous for population 0, heterozygous, and homozygous for population 1, in that order and each 3-dimensional emission probability vector *o*_*i*_ is given by the *i*th neural network output.

While a simple HMM provides a viable approach, it ignores the non-independence of neural network outputs due to overlapping input information and linkage disequilibrium between closely spaced SNPs. To account for this, we model the emission probabilities themselves as arising from a Markov process, assuming that transition probabilities between output pairs vary codependently by class and position. Specifically, we define

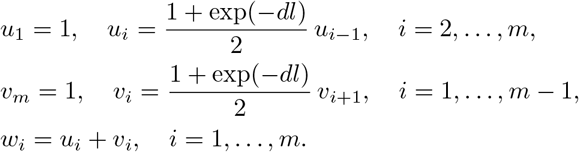

where *m* is the number of outputs. Emission probabilities are then modified as

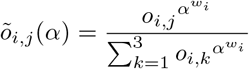

and transition matrices are also adjusted:

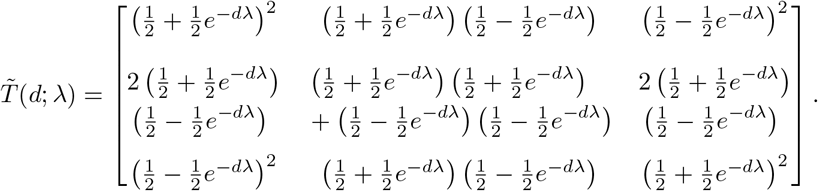

This formulation allows an HMM on the domain *α* ∈ (0, ∞) and *λ* ∈ [0, ∞) to closely approximate the original HMM under the stated assumptions.

We also define a certainty function *c*(*α*) of range (1*/*3, 1) as the mean certainty across all emission vectors, where the certainty of a probability vector is given by its maximum element. For simplicity, we assume that the prior distribution for *α* is such that *c*(*α*) takes on a uniform distribution over its range. We can also calculate the HMM likelihood of *λ* given *α*, as the product of likelihoods of each hidden state given each emission state. The likelihood of an entire solution is further evaluated by its number of switch points, modeled as a Poisson distribution with mean

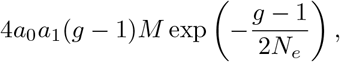

for a chromosome of *M* morgans. This gives us

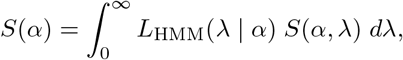

where *S* represents a solution formed by the given HMM parameters. As the likelihood function used here is sharply peaked, *S*(*α*) can be approximated via a maximum likelihood estimate. The solution can then be given by

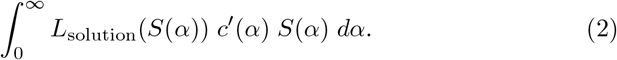

#### 2.2.4 Without reference panels: forming a solution via an iterative approach

When only admixed samples are available, no ancestral information is known initially. We therefore develop an iterative algorithm that converges to a solution. At its core, the algorithm maintains a solution tensor *S*, where *S*_*i,j,k*_ denotes the probability of class *k* for individual *i* at position *j*. At each timestep, a solution vector *S*_*i,j*_ can be updated by treating the remaining samples as reference individuals, using their (probabilistic) classifications as labels. Furthermore, given an output vector *S*_*i,j*_, we can update any other position *b* by

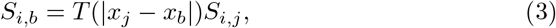

where *T* is the transition matrix defined in Equation (1), and *x* is the vector of genetic positions.

This framework raises two main challenges: (i) how to weight predictions across timesteps, and (ii) how to balance neural network outputs with the transition-based updates from Equation (3). Predictions made early in the algorithm are generally less accurate, since they are based on limited information. Later predictions tend to be more reliable, but giving them too much weight risks neglecting the smoothing effect of earlier outputs and can reduce overall accuracy. A similar tradeoff exists between neural network outputs and transition-based updates. Neural network predictions directly integrate information from neighboring SNPs and individuals, making them highly informative at the focal position. By contrast, the transition updates act as priors that enforce genomic consistency by propagating information across distance. These priors are valuable for smoothing but lose reliability as distance increases, since ancestry states become less correlated. Overweighting them leads to less accurate predictions, while underweighting slows convergence somewhat drastically and reduces smoothing.

To address these issues, we define a credibility score for each neural network output based on the certainty of the labels it uses. The certainty of a label vector *ν* is quantified as

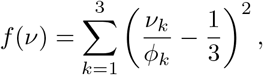

where *ϕ* is the prior class distribution determined by the admixture proportion and is also the stationary distribution of *T*. The credibility of an output at index (*i, j*) is then

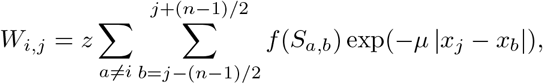

where importance decays exponentially with distance according to parameter *µ*. Here,

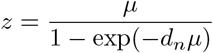

is a normalization term such that the average credibility value does not vary based on the value of *µ*. Odd integer *n* is the number of positions taken by the neural network and *d*_*n*_ is the average distance of any two positions (*n* − 1)*/*2 SNPs apart. This scheme extends naturally to *T*-based updates: the credibility of an update from index (*i, j*) to (*i, c*) is

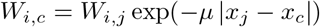

for the same *µ*.

At each timestep, the algorithm stores a tensor *S* of predictions and a matrix *W* of certainties. Since ancestry labels are arbitrary without references, we fix a random individual *a* and position *j* at iteration 0, assume (w.l.o.g.) that it is not homozygous for ancestry 0, and update *S*_*a*_ accordingly. In subsequent iterations, a prediction index (*i, j*) is chosen by weighted random sampling, where the weight is inversely proportional to certainty. A new prediction *s* and credibility *w* are then computed, and the updates

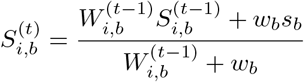

followed by

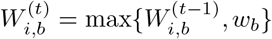

are applied to all positions *b* for timestep *t*. In practice, this update is done in batches and parallelized across multiple individuals.

Since the certainty of each neural network output is assumed to be a linear combination of certainties of other elements of *S*, and since the values of *S* contain predictions across timesteps, with varying amounts of information, it follows that the value of *µ* can be estimated from the predictions themselves. In particular, the optimal value of *µ* is the value that maximizes the sum of certainties of all elements of *S* after a fixed number of updates are done in parallel. This yields a convex optimization problem and can be solved via golden-section search. The value of *µ* is not assumed to be constant throughout the algorithm; it is updated periodically as the algorithm progresses.

The iterative nature of this algorithm makes it prone to convergence at local optima, a problem that becomes more frequent as switch points increase. Some non-optimal solutions can be detected statistically, but others are subtler. To mitigate this, the genome is partitioned into overlapping windows, each run with multiple random seeds. After convergence, a global solution is assembled by selecting the sequence of windows that maximizes similarity across overlaps, while accounting for the fact that ancestry references are local to each window. In addition to detecting examples of bad convergence, this approach both smooths predictions and resolves inconsistencies in local ancestry references.

In practice, this process is implemented dynamically. Evaluation metrics identify problematic regions, which are recursively subdivided into smaller windows (each 2*/*3 of the parent size). Consecutive triplets of windows are tested to ensure consistency of ancestry labels across regions. This refinement continues until a predefined total number of windows is reached.

#### 2.2.5 Handling unknown parameters

Our method requires two parameters—admixture proportion and admixture time—which may or may not be known in advance.

Admixture proportion can be estimated directly from the neural network outputs. At the start of the algorithm, we run a round of neural network inference and use the raw class probabilities to estimate the global ancestry proportion. When multiple admixed samples are available, their estimates can be combined to obtain a more stable proportion before running the full algorithm.

Admixture time is more challenging to infer. We begin with a prior distribution, either specified by the user or taken to be uniform over 1–1000 generations in the absence of prior information. Using this distribution, we run the full inference algorithm, substituting the prior whenever admixture time is needed. From the resulting distribution of solutions (Equation (2)), we derive a corresponding distribution of switch point counts. The likelihood of observing *s* switch points given *g* generations since admixture is modeled as

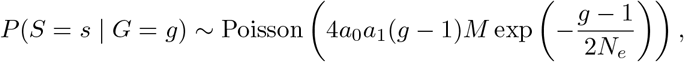

and *P* (*G* = *g* | *S* = *s*) can be calculated using Bayes’ rule. Our updated distribution for admixture time then becomes

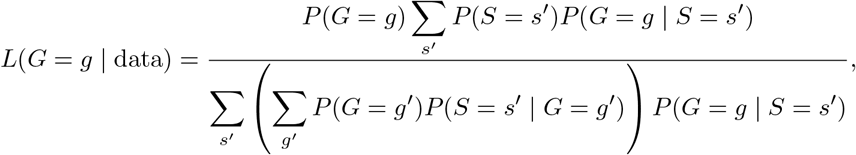

where the denominator corrects for non-reciprocity between distributions of *S* and *G*. This updated distribution becomes the prior for the next iteration. When multiple admixed samples are available, the process cycles through each sample, updating the distribution after every iteration.

The final output consists of both the posterior distribution for admixture time and the inferred local ancestry along the chromosome. For now, this procedure is only implemented in the reference-panel setting, although a similar scheme could be adapted to the panel-free case.

## 3 Results

We evaluate our method across three datasets: human, *Drosophila melanogaster*, and fully simulated populations generated using MACS. For each dataset, we benchmark all methods under a range of admixture proportions and admixture times. In each evaluation, we measure diploid accuracy on a single admixed individual using 16 reference individuals from each of two ancestries. For our reference-free variant, we instead test accuracy on the same admixed individual using 32 additional admixed samples of unknown ancestry. Switch points for admixed chromosomes are generated using SELAM [13], and computational phasing is performed with Beagle [14] where required. We compare our method against RFMix [2] and LOTER [15], two widely used local ancestry inference methods that have demonstrated strong performance with modest reference panel sizes.

### 3.1 Human Dataset

For the human dataset, we use European (CEU) and Chinese (CHB) populations from the 1000 Genomes Project [16]. ORIAN achieves the highest median diploid accuracy for admixture times below 32 generations, after which performance declines more rapidly compared to other methods. When run without reference panels, accuracy did not reach biologically meaningful levels, and we do not report these results here.

**Figure 1.**
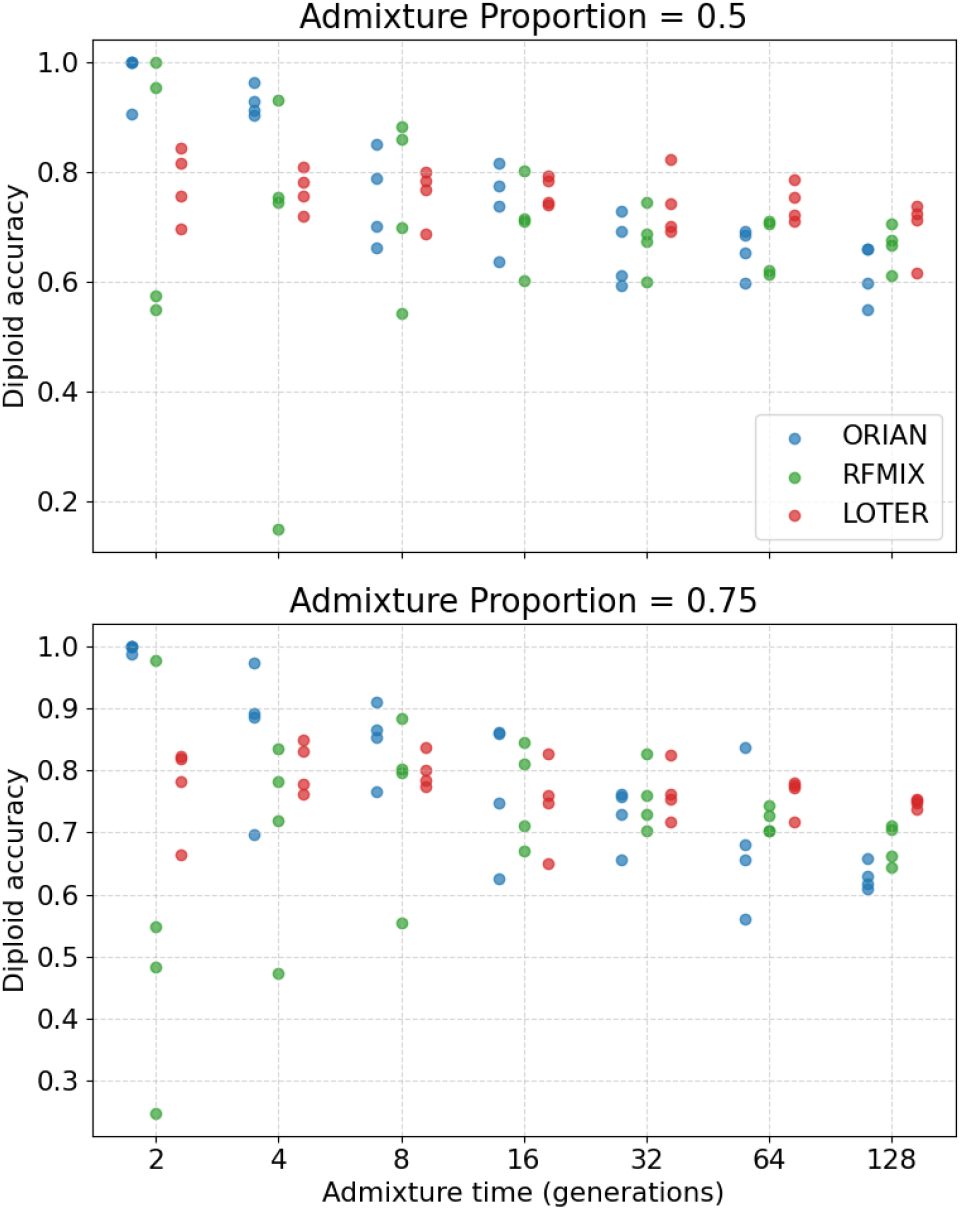
Diploid accuracy of ORIAN, RFMIX, and LOTER for simulated admixture between European (CEU) and Chinese (CHB) populations. Reference and admixed haplotypes are sampled from the 1000 Genomes Project. Computational phasing is performed with Beagle, using only information available within each simulated scenario.

### 3.2 *Drosophila melanogaster* Dataset

To evaluate performance in a different genomic context, we simulate admixture between European and African populations of *D. melanogaster*, using genomic data obtained from FlyBase [17]. Here, ORIAN outperforms both RFMIX and LOTER across all parameter settings. Notably, when using only other admixed individuals for local ancestry inference, ORIAN’s median diploid accuracy continues to exceed that of competing methods for values of admixture time up to and including 64 generations, despite the increased difficulty of the task.

**Figure 2.**
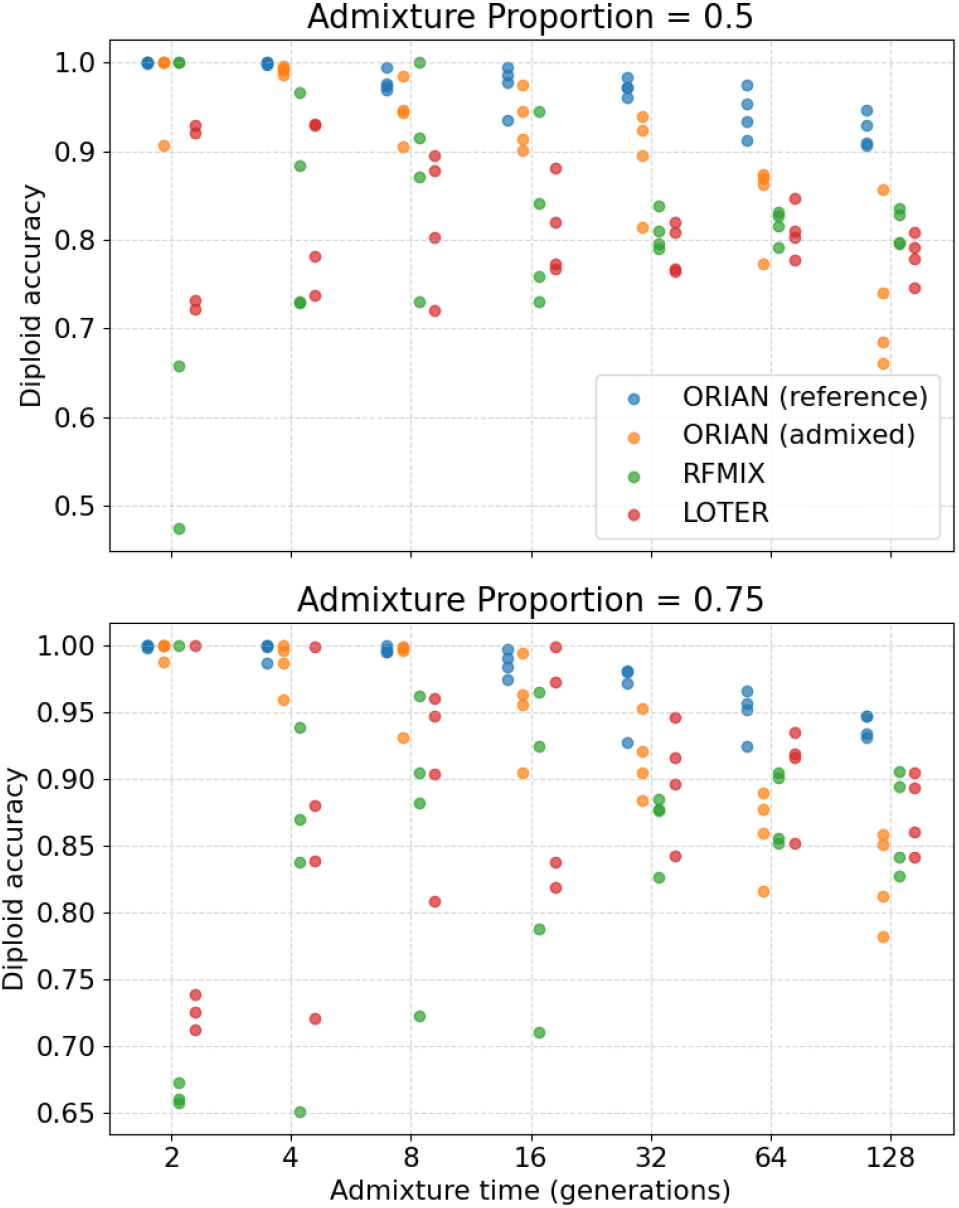
Diploid accuracy of ORIAN, RFMIX, and LOTER for simulated admixture between European and African populations of *D. melanogaster*. Reference and admixed haplotypes are obtained from the Flybase dataset. Computational phasing is performed with Beagle using only information available within each simulated scenario.

### 3.3 Fully Simulated Populations

Finally, we evaluate ORIAN on a dataset of fully simulated populations, generated using MACS [10] to model ancestral divergence. Simulation parameters were sampled randomly and independently to capture a broad range of biologically plausible scenarios. In this setting, ORIAN consistently outperforms both RFMIX and LOTER when reference individuals are available. When relying solely on other admixed individuals, ORIAN’s accuracy is slightly lower but remains comparable to that of the other methods, despite using substantially less information.

**Figure 3.**
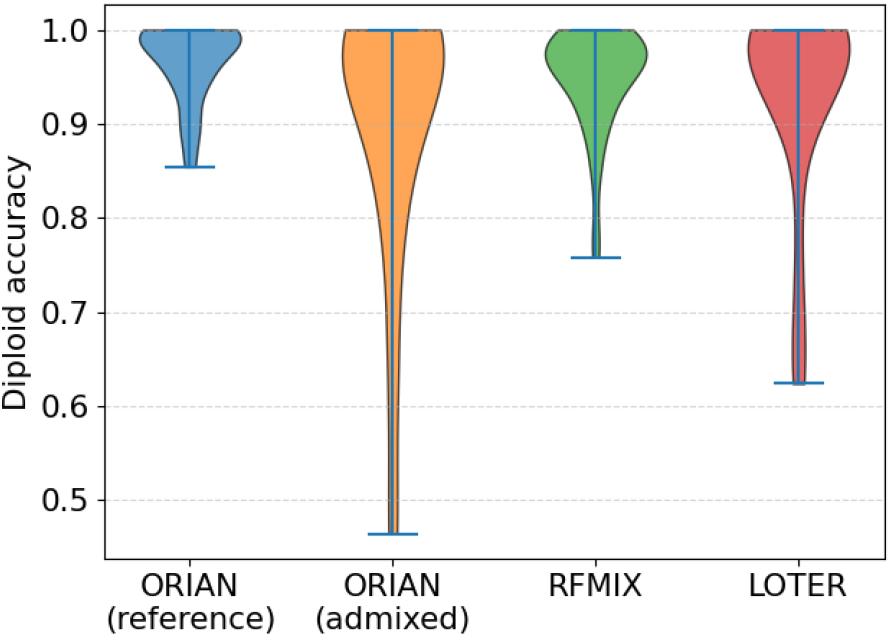
Diploid accuracy of ORIAN, RFMIX, and LOTER for simulated admixture between two populations generated with MACS and SELAM. Parameters are drawn randomly from biologically plausible ranges. Results are shown as violin plots to illustrate both accuracy and variability across simulations.

### 3.4 Computational Performance

To evaluate computational efficiency, we measured wall-clock runtime for ORIAN, RFMix, and LOTER under the same experimental conditions used in the accuracy benchmarks. All methods were run on the same hardware environment, and runtimes were averaged across multiple runs to reduce variability. For ORIAN, we report results using GPU acceleration, while RFMix and LOTER were run on CPU, as these methods do not natively support GPU-based inference. Each reference-based evaluation consists of inferring local ancestry along a single admixed chromosome using 16 reference individuals per population, whereas the reference-free setting involves jointly inferring the ancestry of 33 admixed individuals.

**Table 1.** Comparison of wall-clock runtime (in seconds) across methods and datasets. Values are averaged over multiple runs under identical experimental conditions. Runtime for ORIAN (reference-free) corresponds to joint inference across 33 admixed individuals, with average runtime per individual in parenthesis, whereas other methods are applied to a single individual. Phasing time is added to total runtime of relevant methods.

| Method | Hardware | Human | *D. mel* | Simulated |
| --- | --- | --- | --- | --- |
| ORIAN (ref-based) | GPU | 291.8 | 280.9 | 297.2 |
| ORIAN (ref-free) | GPU | N/A | 1567.0 (47.5) | 1181.8 (35.8) |
| RFMix | CPU | 76.3 | 32.9 | 70.8 |
| LOTER | CPU | 47.4 | 13.1 | 39.9 |

Across all datasets, ORIAN requires more computation time than both RFMix and LOTER, even when implemented with parallelization and hardware acceleration. This reflects the increased model complexity, including neural network inference, multiple forward passes, and, in the reference-free setting, an iterative procedure that updates ancestry estimates over multiple timesteps.

In the reference-free setting, runtime depends on the number of generations since admixture, as the expected number of ancestry switch points grows with this parameter. Because the iterative algorithm must resolve these switch points across individuals, the total number of updates increases accordingly, resulting in approximately linear scaling of runtime with *g*. To capture this behavior, we additionally evaluate runtime as a function of admixture time using the *D. melanogaster* dataset. As expected, ORIAN is substantially faster for recent admixture scenarios (small *g*), while runtime increases for more ancient admixture events.

**Table 2.** Runtime (in seconds) of ORIAN (reference-free) as a function of generations since admixture. Values are averaged across eight LAI scenarios for each admixture time.

| Admixture time ($g$) | Total runtime | Runtime per individual |
| --- | --- | --- |
| 2 | 435.2 | 13.2 |
| 4 | 465.4 | 14.1 |
| 8 | 510.6 | 15.5 |
| 16 | 971.5 | 29.4 |
| 32 | 2147.4 | 65.1 |
| 64 | 2895.9 | 87.8 |
| 128 | 3543.3 | 107.4 |

Despite this increased computational cost, ORIAN provides several important advantages. In the reference-based setting, it achieves improved accuracy in scenarios with limited reference panel sizes. More importantly, ORIAN enables local ancestry inference without access to reference panels, a setting in which existing methods such as RFMix and LOTER cannot be directly applied. These results highlight a tradeoff between computational efficiency and modeling flexibility, with ORIAN offering improved performance in challenging settings at the expense of increased runtime. In many applications, inference is performed once per dataset, making runtime a secondary consideration relative to accuracy and applicability.

## Acknowledgments

The author thanks Professor Russell Corbett-Detig for proposing the original research direction and providing guidance on the project. This work was initiated while the author was employed as an Academic Researcher at the University of California, Santa Cruz.

## Code Availability

The ORIAN source code is available at: https://github.com/thomasjdiem/ORIAN

